# Phylogenomic insights into the first multicellular streptophyte

**DOI:** 10.1101/2023.11.01.564981

**Authors:** Maaike J. Bierenbroodspot, Tatyana Darienko, Sophie de Vries, Janine M.R. Fürst-Jansen, Henrik Buschmann, Thomas Pröschold, Iker Irisarri, Jan de Vries

**Affiliations:** University of Goettingen, Institute for Microbiology and Genetics, Department of Applied Bioinformatics, Goldschmidtstr. 1, 37077 Goettingen, Germany; University of Goettingen, Campus Institute Data Science (CIDAS), Goldschmidstr. 1, 37077 Goettingen, Germany; University of Applied Sciences Mittweida, Faculty of Applied Computer Sciences and Biosciences, Section Biotechnology and Chemistry, Molecular Biotechnology, Technikumplatz 17, 09648 Mittweida, Germany; University of Innsbruck, Research Department for Limnology, 5310 Mondsee, Austria; Section Phylogenomics, Centre for Molecular Biodiversity Research, Leibniz Institute for the Analysis of Biodiversity Change (LIB), Museum of Nature, Hamburg, Martin-Luther-King Platz 3, 20146 Hamburg, Germany; University of Goettingen, Goettingen Center for Molecular Biosciences (GZMB), Department of Applied Bioinformatics, Goldschmidtstr. 1, 37077 Goettingen, Germany

**Keywords:** Charophyta, streptophyte algae, multicellularity, phylogenomics, plant terrestrialization, plant evolution, ancestral character state

## Abstract

Streptophytes are best known as the clade containing the teeming diversity of embryophytes (land plants)^1–4^. Next to embryophytes are however a range of freshwater and terrestrial algae that bear important information on the emergence of key traits of land plants. Among these, the Klebsormidiophyceae stand out. Thriving in diverse environments—from mundane (ubiquitous occurrence on tree barks and rocks) to extreme (from the Atacama Desert to the Antarctic); Klebsormidiophyceae can exhibit filamentous body plans and display remarkable resilience as colonizers of terrestrial habitats^5,6^. Currently, the lack of a robust phylogenetic framework for the Klebsormidiophyceae hampers our understanding of the evolutionary history of these key traits. Here, we conducted a phylogenomic analysis utilizing advanced models that can counteract systematic biases. We sequenced 24 new transcriptomes of Klebsormidiophyceae and combined them with 14 previously published genomic and transcriptomic datasets. Using phylogenomic analysis built on 420 loci and sophisticated models, we establish a novel phylogenetic structure, dividing the six distinct genera of Klebsormidiophyceae in a novel four-order-system, with deep divergences more than 898, 765, and 734 million years ago. The reconstruction of ancestral states for habitat suggests an evolutionary history of multiple transitions between terrestrial-aquatic habitats, with Klebsormidiales having conquered land earlier than embryophytes. Focusing on the body plan of the last common ancestor of Klebsormidiophyceae, we postulate it was likely filamentous whereas the sarcinoids and unicells in Klebsormidiophyceae are likely derived states. Our data reveal that the first multicellular streptophytes likely lived more than 900 million years ago.

## RESULTS AND DISCUSSION

### Klebsormdiophyceae are cosmopolitan colonizers of diverse habitats

In our sampling, we accounted for the class of Klebsormdiophyceae standing out among green algae by their resilience^6,7^ and habitat range. We obtained representatives that can be found in streams, rivers^8,9^, lakeshores^10^, bogs^9^, soil^11^, natural rocks in flat and mountainous regions^12^, tree bark^13^, acidic post-mining sites, freshwater bodies^14–16^, sand dunes^17^, biotic crusts of hot deserts^18^, and human-shaped habitats such as urban walls^19^, and building façades^20,21^. We sampled from the hottest (Atacama Desert) to the coldest (Antarctic) regions, from growing in fresh water to land, including representatives involved in forming biological soil crusts (BSCs). Overall, our sampling presents a worldwide distribution map for the ancient lineage of Klebsomidiophyceae, showcasing their habitat utilization and adaptability, ecological significance, and hidden diversity (Figure 1A and B).

**Figure 1:**
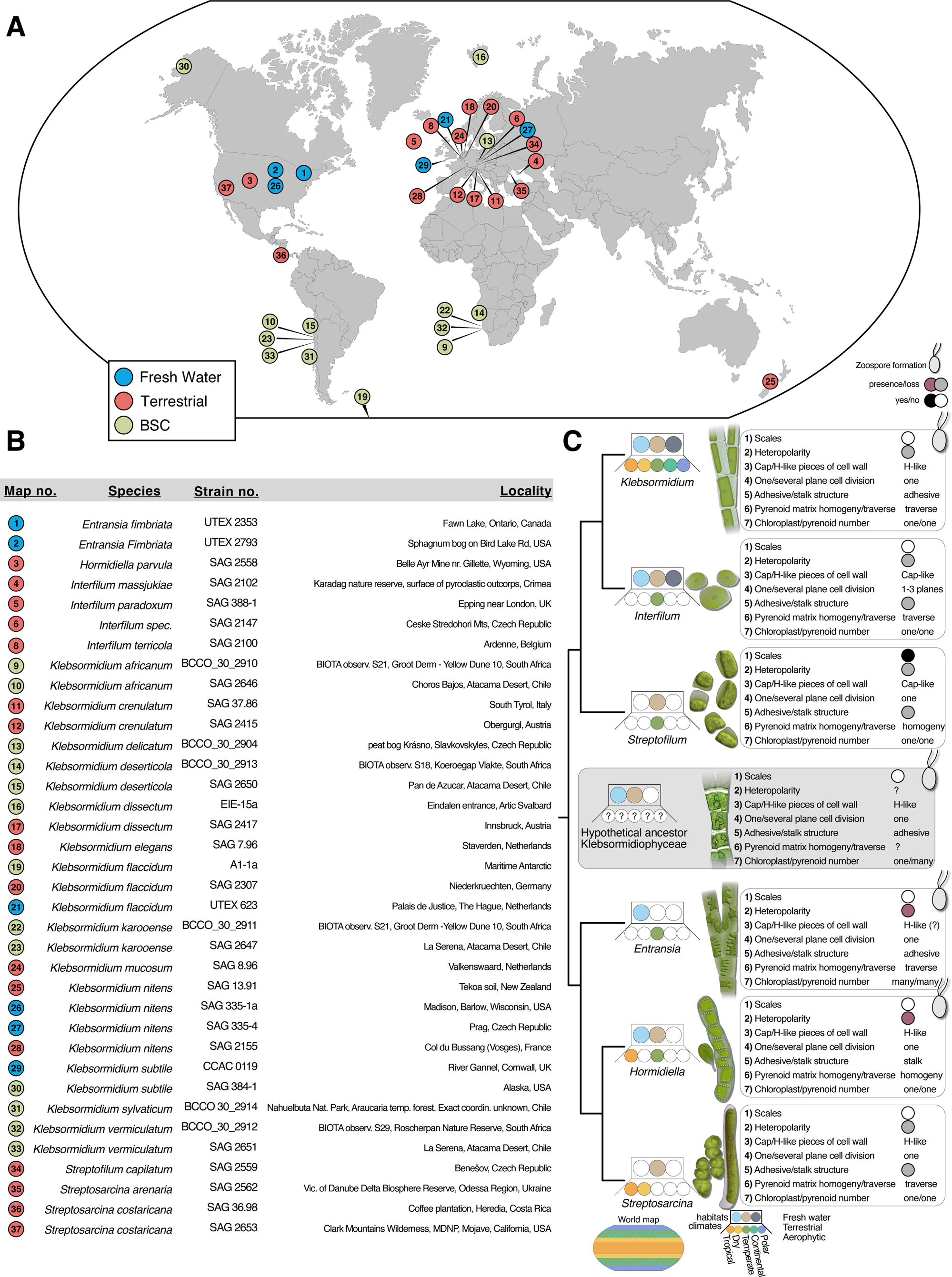
Biogeography of Klebsormidiophyceae. **A:** World map with all the klebsormidiophycean strains used within this study. An interactive map can be accessed under https://tinyurl.com/yph2s4ma. **B:** Details on the strains of Klebsormidiophyceae used in this study. **C**: Cladogram of the genera in Klebsormidiophyceae. Dots label their distribution across climate zones, habitats, and body plan diversity. Character information were guided by Mikhailyuk et al.^26^

The ability of Klebsormidiophyceae to dwell in diverse habitats is underpinned by a set of molecular physiological traits, such as those contributing to desiccation resistance. For example, *Klebsormidium crenulatum* can undergo regulated cell shrinkage under desiccation stress supported by a flexibility of its cell wall and the accompanying alterations in the actin cytoskeleton^6^. Furthermore, in terrestrial habitats, temperature fluctuations are more pronounced compared to aquatic environments. As a result, membrane fluidity becomes crucial, as it directly impacts the expression of genes involved in cold or heat responses^22–24^. Owing to the broad distribution of habitat adaptations (Figure 1C), the emergence of the enabling traits needs to be projected onto a phylogenetic framework.

The class Klebsormidiophyceae comprises a minimum of five different genera: *Klebsormidium, Interfilum, Entransia, Hormidiella, and Streptosarcina*^25,26^. Within *Interfilum* and *Klebsormidium*, seven main clades (A, B, C, D, E, F, and G) have been identified through the analysis of ITS and *rbc*L markers^21,27,28^. To scrutinize the distribution and diversity of Klebsormidiophyceae, we studied 24 strains and included 14 previously published isolates in this dataset; we gave particular attention to sample densely within clade G, which is rich in species but currently scarce in sequence data. Preliminary phylogenetic analyses were performed using a congruent dataset of 31 streptophycean taxa, employing two commonly used markers (*rbc*L, SSU; Supplementary Figure 1). Owing to the incongruence between the phylogenetic trees built on marker genes, no robust reconstruction of the relationships within Klebsormidiophyceae was possible (Supplementary Figure 1). Indeed, in line with previous studies, two major problems were unsolved: (i) The lack of monophyly of *Klebsormidium*. The type species of *Klebsormidium* (*K. flaccidum*; clades B/C) represents the sister to *Interfilum* based on *rbc*L and ITS sequences, whereas the rest of *Klebsormidium* (clades D-G) remains separated from those. If true, this had crucial taxonomical consequences because of the priority rule: the genus *Interfilum* is older than *Klebsormidium* and would have priority if these phylogenetic results were robust. (ii) The phylogenetic position of the genus *Streptofilum* within the Streptophyta. According to the phylogenetic analyses of the combined *rbc*L-SSU and 44 chloroplast genes, *Streptofilum* is suggested to represent a separate lineage outside of Klebsormidiophyceae^26,29^, however, the phylogeny remains unresolved. Overall, the resolution of the marker genes is not powerful enough to resolve the complex evolutionary history of the Klebsormidiophyceae. To scrutinize the internal genetic structure of Klebsormidiophyceae, we took advantage of the higher resolving power of phylogenomics based on hundreds of genes.

### A phylogenomic framework and four-order-system for Klebsormidiophyceae

Using the Illumina NovaSeq6000 platform, we sequenced the transcriptomes of 24 isolates of Klebsormidiophyceae, including 15 strains of *Klebsormidium*, 4 *Interfilum*, and one of each *Hormidiella* and Streptofilum as well as three isolates of *Streptosarcina* collected from five continents and various habitats from all climate zones. In total, we sequenced 1.407 billion paired-end transcriptomic reads, providing more than 423 gigabases of raw sequence information. To complement our dataset, we integrated these data with 14 previously published transcriptomes and 24 additional samples of algae and land plants (see Methods). With a focus on 420 densely sampled loci, we used maximum likelihood with the complex LG+F+I+Γ4+C60 mixture model to construct a robust phylogenomic tree (Figure 2).

**Figure 2:**
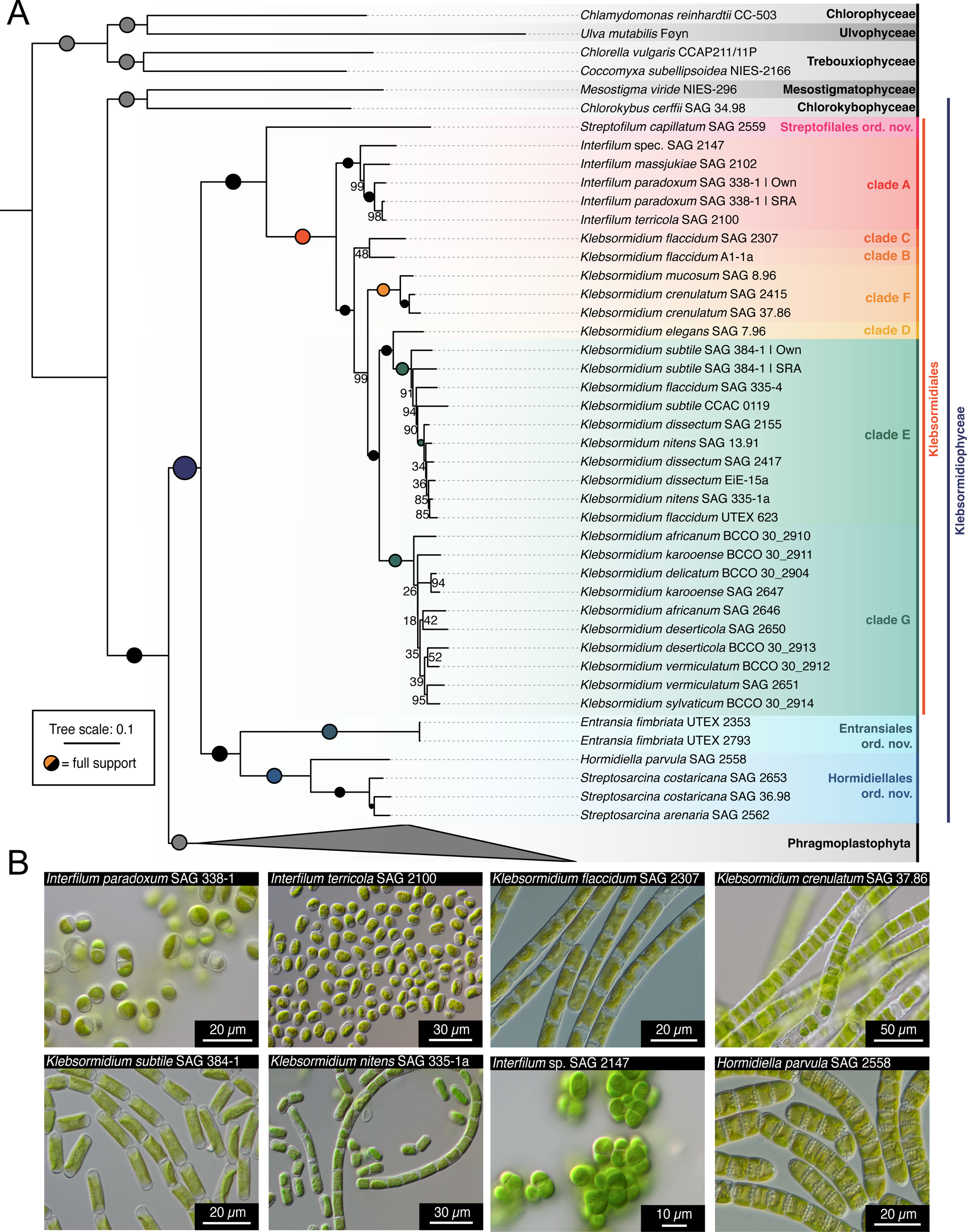
A new four-order-system of Klebsormidiophyceae based on phylogenomics. **A:** Maximum likelihood phylogenetic analyses based on 420 loci and the complex LG+F+I+Γ4+C60 mixture model. An SH-like aLRT branch support test was employed and the size of the dots corresponds to the support values (legend). **B:** A selection of the morphological diversity found across Klebsormidiophyceae.

The maximum likelihood tree was fully resolved and accurately represented the accepted phylogeny of the green lineage (Chloroplastida). The tree confirmed the positioning of the Klebsormidiophyceae as the sister group to all Phragmoplastophyta^1,30^ with full non-parametric bootstrap support (Figure 2). Within Klebsormidiophyceae, two major clades emerged: (i) a clade consisting of *Entransia*, *Hormidiella*, and a monophylum of the three *Streptosarcina* strains that branched sister to (ii) the clade of the other Klebsormidiophyceae, consisting of clades A to G (formed by the genera *Interfilum* and *Klebsormidium*) and *Streptofilum*. Importantly, we recovered a monophyletic genus *Klebsormidium* (25 strains, full support) and a monophyletic genus *Interfilum* (4 strains, full support; clade A). *Klebsormidium flaccidum* (SAG 2307) and *Klebsormidium flaccidum* (A1-1a) formed an assemblage (a non-supported monophylum) next to the other *Klebsormidium* spp., yielding a clades B and C^21^. Indeed, while the genus *Klebsormidium* was monophyletic, some species were recovered as paraphyletic and will require taxonomic revision. Thus, *Klebsormidium* and *Interfilum* form a monophylum, to which *Streptofilum capillatum* branches as sister.

Our phylogenomic analyses recover a topology that features a deep split in the Klebsormidiophyceae. According to our molecular clock analyses, this split happened 898.46 (702.56-1147.5 95% HPD age estimates) million years ago (Suppl. Figure 2). The resulting two clades again split 734.2 (565.56-945.55 95% HPD age estimates; *Streptofilum*–*Interfilum*/*Klebsormidium*) and 764.97 (591.72-977.45 95% HPD age estimates; *Entransia*–*Hormidiella*/*Streptosarcina*) million years ago. We recover that even these two more shallow splits are deeper than the deepest divergence in Embryophyta (for timing on embryophytes, see Morris et al.^31^). We therefore propose to account for this deep genetic structure by diving Klebsormidiophyceae into a four-order system (Table 1): Klebsormidiales with the genera *Interfilum* and *Klebsormidium* (forming a fully-supported monophylum that encompasses the clades and grades A to G), Hormidiellales ord. nov., Streptofilales ord. nov., and Entransiales ord. nov. (Figure 2).

**Table 1:**
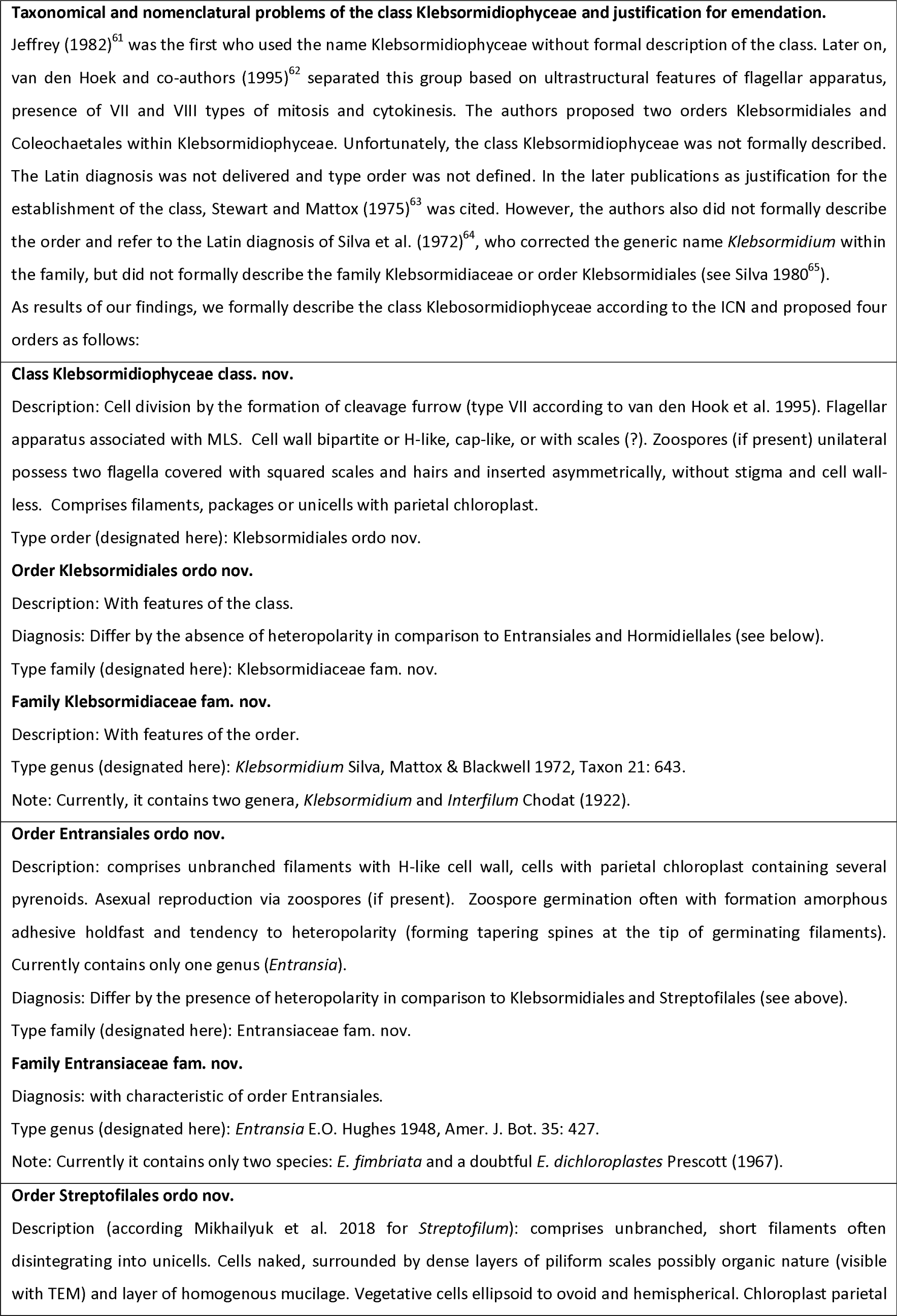
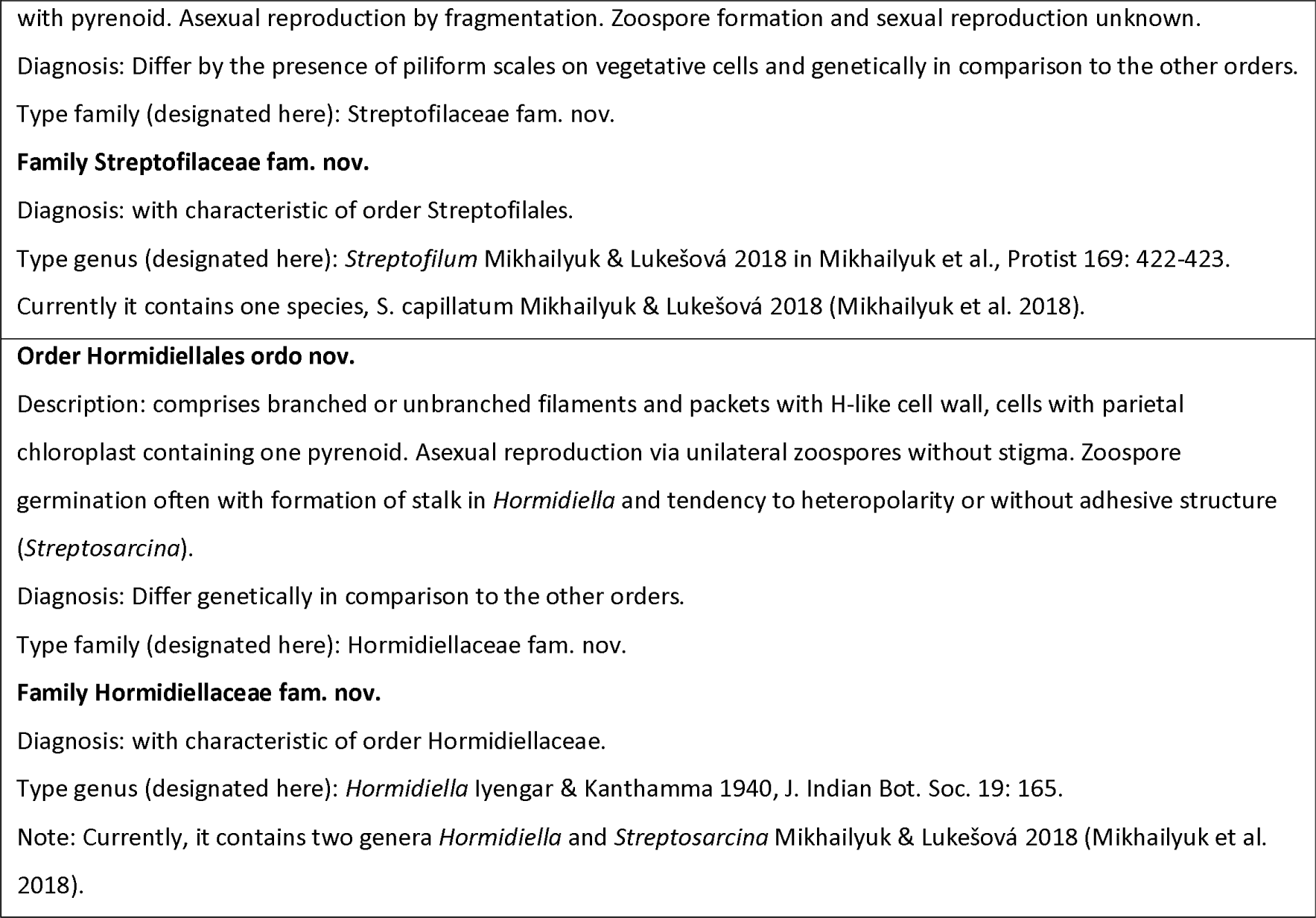
Revision of the class Klebsormidiophyceae and its orders.

The enigmatic *Entransia* was first described from Nova Scotia^32^ and for a long time tentatively placed in the Zygnematophyceae. We here describe it as Entransiales ord. nov., consisting of a fully-supported clade and the family Entransiaceae fam. nov., forming a clade with the Hormidiellales ord. nov. Previous molecular phylogenetic analyses based on several genes demonstrated that *Entransia* is a member of Klebsormidiophyceae^33–35^. Cook^36^ conducted a detailed study of the two available isolates of *Entransia* and firmly anchored its position within Klebsormidiophyceae solely based on morphological and cytological features. The morphological characters shared by *Entransia* and other Klebsormidiophyceae include cylindrical cells, unbranched filaments, parietal laminated chloroplast, H-shaped cross-wall pieces, asexual reproduction by fragmentation as well as zoospores and aplanospores.

*Hormidiella* is here described as part of Hormidiellales ord. nov. and the family Hormidiellaceae fam. nov.. Both *Entransia* to *Hormidiella* have the tendency for upright growth and differentiation into three types of vegetative cells: cells with adhesive structures (*Entransia*) or stalks (*Hormidiella*), normal vegetative cells and upper tapered cell. Both genera also produce asexual zoospores. The sister genus *Streptosarcina* appears to have lost asexual zoospores during the adaptation to the arid habitats and developed instead 2D-division^26^, which may serve as protection from desiccation; the mechanisms and induction of branching in *Streptosarcina* however remain obscure. Some traits are also shared by G-clade *Klebsormidium* spp. and *Hormidiella*. Morphological investigation of mature cultures demonstrated that these organisms possess relative coin-like cells and filaments disintegrating into short filaments^26^ with tapered end cells. Representatives of the G-clade have generally smaller cell sizes that could also be adaptative to desiccation.

*Streptofilum* is here described as part of Streptofilales ord. nov. and the family Streptofilaceae fam. nov., which are sister to the clade of *Klebsormidium* and *Interfilum* (the Klebsormidiales). In the description of *Streptofilum*, the authors^26^ noted a shared feature with mature vegetative cells of *Mesostigma*: a scaly cell wall. Other representatives of the Streptophyta, including some mosses and ferns, possess cell-covering scales only in asexual reproductive stages (zoospores). Hence, while Streptofilum and Interfilum might have superficial similarities in morphology and ecology, the ultrastructure of the vegetative cells showed organic scales in *Streptofilum* (in contrast to *Interfilum*). This is underscored by the 734-million-year divergence between both genera.

Overall, while there is a set of shared and distinct traits to all orders in Klebsormidiophyceae, our phylogenomic data establish a robust and unequivocal backbone of Klebsormidiophyceae evolution consisting of two deep dichotomies nested within an even deeper dichotomy in Klebsormidiophyceae— each of these splits being more distant in the past than the split of all land plants. We next used this new phylogenomic framework and four-order-system to understand the evolutionary history of key traits.

### The first multicellular streptophyte emerged more than 900 million years ago

Land plants are among those photosynthetic eukaryotes with the most complex true multicellularity. The evolutionary emergence of streptophyte multicellularity is thus one of the general interest topics. Here, streptophyte algae hold important information and surprises. Among those streptophyte algae most closely related to land plants, the Zygnematophyceae, we find unicells and (at times branched) filaments. This stands in stark contrast to other (phragmoplastophytic) streptophyte algae, which have parenchymatous growth (Coleochaetophyceae) or even erect growth with organs and thus 3D growth^37^ (Charophyceae). It was inferred that the common ancestor of Zygnematophyceae likely underwent reduction and might have even ancestrally transitioning to a unicellular body^38^. To understand the propensity for multicellularity among Phragmoplastophyta, one must turn to its sister group: the Klebsormidiophyceae.

Klebsormidiophyceae includes sarcinoid (a thallus comprised of cellular colonies organized in a three-dimensional, packet-like structure), uniserial unbranched filaments, and filaments that easily disintegrate into unicells^26,28^. According to phylogenetic reconstruction based on single (or few) gene(s) or multigene approaches^21,25,26^, the position of the sarcinoid morphotype is spread among different clades and probably is derived from filamentous type. Interestingly, both sarcinoid and filamentous morphotypes are present sometimes within the same species or genus, for example in *Streptosarcina costaricana* or *Interfilum*^25,26^. This could represent an advantage for colonizing different terrestrial substrates, due to the lower surface-to-volume relationship of large cell assemblages or the possibility of crust formation by filaments. To understand the evolution of growth types in Klebormidiophyceae and streptophytes in general, we conducted Ancestral Character State Reconstructions (ACSR) by maximum likelihood. Multiple data coding strategies were employed, particularly focused on the type of cellular growth (Figure 3; Supp. Figure 3). In the simplest coding pattern, we recovered full support for a multicellular ancestor of Klebsormidiphyceae (posterior probability [PP] of 0.992) and for a multicellular ancestor of Klebsormidophyceae and Phragmoplastophyta (PP of 0.988). If we employ a three-character coding, distinguishing between unicells, sarcinoid cell packages, and filamentous or more complex body plans, we also recover a multicellular ancestor of Klebsormidophyceae and Phragmoplastophyta, with a likely filamentous (PP of 0.620 and 0.613) and less likely sarcinoid (PP of 0.359 and 0.366) body plan. Thus, the first multicellular streptophyte likely lived around 900 million years ago.

**Figure 3:**
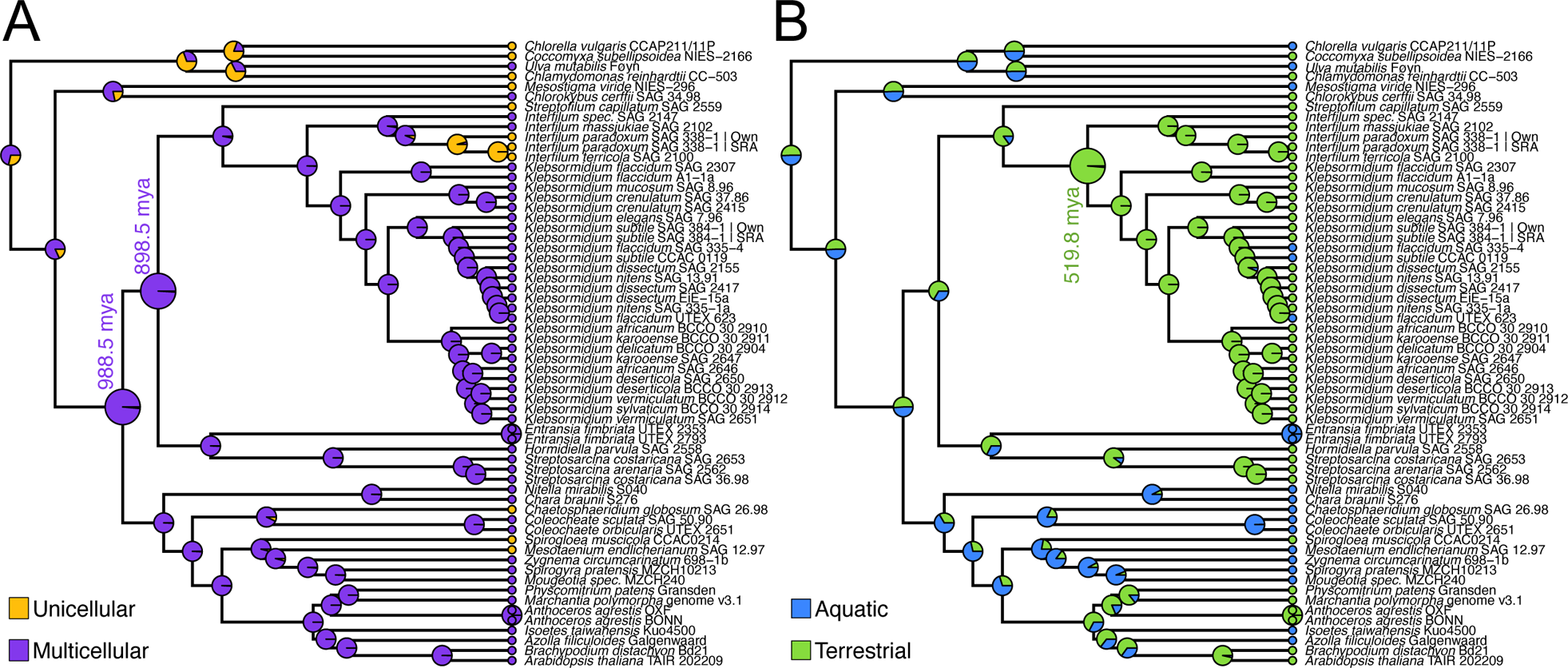
Ancestral character state reconstruction of body plan and habitat characters across 900 million years of klebsormidiophyceaen evolution. **A**: To examine the ancestral character states of growth types in unicellular or multicellular organisms, coding schemes represented varying levels of complexity and hypotheses regarding the homology of growth types. The shown color-coded character state distributions represents yellow for unicellular growth, purple for multicellular growth sensu lato (including sarcinoid, filamentous, and parenchymatous growth); a different coding scheme is shown in Supp. Figure S3. Note ancestor nodes for (i) Klebsormidiophyceae and Phragmoplastophyta 988.5 mya and (ii) Klebsormidiophyceae 898.5 mya. **B**: To examine the ancestral habitats of the Klebsormidiophyceae, we coded the habitat occurrence of the species as blue for aquatic and green for terrestrial. Note the terrestrial ancestor of Klebsormidiales, which lived 290.4 mya; a different coding scheme is shown in Supp. Figure S3. Divergence dating is based on the molecular clock analyses shown in Suppl. Figure S2.

What does this mean for the ability to grow on land? While some streptophyte lineages, such as *Mesostigma* or *Chlorokybus*, are rare algae with very narrow ecological niches, one of the most striking successes in colonizing terrestrial habitats can be found in *Klebsormidium*. *Klebsormidium* spp. are one of the few eukaryotes that are capable of forming biological soil crusts on their own or together with cyanobacteria, mosses, or lichens^39^. We coded habitat occurrence and, using our newly established phylogenetic backbone, performed ancestral character state reconstruction to identify the history of habitat shifts in Klebsormidiophyceae. No clear pattern support was recovered for the deep ancestors, but a full support (PP of 0.994) for ancestrally terrestrial Klebsormidiales (Figure 3B, node 519.84 mya [384.37-692.68 95% HPD age estimates]). This aligns with physiological traits. *Interfilum* and *Klebsormidium* are known to produce special mycosporine-like amino acids acting as UV-protectors^40^. Another source of protectants could be phenylpropanoid-derived compounds. The first enzyme in the pathway, phenylalanine-ammonia lyase (PAL), was thought to be a land plant innovation that was proposed to have emerged via horizontal gene transfer from soil-associated bacteria^41^. However, when the genome of *K. nitens* was published^42^, it was found to have a PAL homolog ^43,44^, raising the question of when the streptophyte PAL emerged. In our dataset, we found candidates for PAL for other Klebsormidiales (*Klebsormidium* and *Interfilum*), which formed a fully-supported clade with bacterial PALs (Suppl. Figure 4). No homologs outside of Klebsormidiales—neither other Klebsormidiophyceae nor other algae—were found. Overall, this suggests, that the ancestor of *Klebsormidium* and *Interfilum* may have acquired a bacterial PAL from soil-associated bacteria independent from the origin in land plants^41^.

Our data suggest that the ability for filamentous growth is ancient in streptophytes — it emerged at least 900 million years ago. Recently, Hess et al.^38^ found that the Zygnematophyceae (the algal sister lineage of land plants) might have (re-)gained a filamentous body plan multiple times independently. Our ACSR on Klebsormidiophyceae might help explain this: The molecular machinery for filamentous growth might be a set of homologous genes shared since the last common ancestor of Klebsormidiophyceae and Phragmoplastophyta. Hence, there is an ancient 900-million-year-old genetic potential for multicellularity among streptophytes, explaining the propensity to become multicellular. This propensity, building on this genetic potential, was realized multiple times throughout streptophyte evolution, sometimes resulting in very complex multicellular organisms like *Chara* and land plants, other times in manifesting in mere filamentous growth. That there is a smooth transition between unicellular and multicellular growth is palpable when taking a closer look at, for example, *Interfilum* sp. SAG2147 whose cells are often organized in clumps (Supplementary Figure 5). However, these clumps easily fall apart, resulting mainly in groups of two or four cells. Groups with four cells can be arranged longitudinally, suggesting that cytokinesis was transverse only. However, in packs of four, it appears that the cell division plane has changed by 90 degrees since the previous division. This indicates rotations in the cell division plane. Additionally, cell wall remodeling after cell division seems to be extensive and fast.

Filamentous Klebsormidiophyceae exhibit, like filamentous Zygnematophyceae, one of the simplest forms of multicellularity: non-branching (“1D”) filaments^37,45^. A classical vegetative centripetal cleavage gives rise to these filaments^46,47^. That said, some intricate molecular mechanisms known from land plants might be at play, foremost the ancient phytohormone auxin^48^. Ohtaka et al.^49^ found that *Klebsormidium nitens* NIES-2285 alters its cell division and cell elongation upon auxin treatment and it was later confirmed that *K. flaccidum* has a functional auxin efflux carrier^50^—which in land plants is key for polar auxin transport-mediated morphogenesis^51^. Thus, some key morphogenic processes likely had a deep evolutionary origin in the first filamentous streptophyte.

The frequent loss and gain of filamentous growth suggest that this habit involves several independent genes, each with additional functions. Indeed, also unicellular green algae have most of the genes needed for multicellularity^52,53^. This implies that the complete loss of these genes, even when the lineage reverts to a simpler growth type (likely unicells), is unlikely. This enables both forward and backward evolutionary transitions in body plans across many clades and millennia, as only one or a few genes need to undergo slight changes in their activity.

## Conclusion

Significant efforts have been made in the past decade to understand the phylogenetic relationships within streptophytes, particularly in relation to embryophytes, and the deep evolutionary origin of land plant traits^30,38,42,54–60^. The evolutionary history of one of the defining traits of land plants however remains debated: multicellularity and complex body plans. We investigate Klebsormidiophyceae using a phylotranscriptomic approach building on isolates from around the world to establish the deep genetic structure within Klebsormidiophyceae and their relation to Phragmoplastophyta. Through ancestral character state reconstruction, we demonstrate that the common ancestor of Phragmoplastophyta and Klebsormidiophyceae was already a multicellular alga; this alga lived almost a billion years ago.

## ACKNOWLEDGEMENTS

This work was funded by the German Research Foundation grant 509535047 (VR 132/10-1) to J.d.V. and the grants 440231723 (VR 132/4-1), 509535047 (VR 132/10-1) to J.d.V. and 440540015 (BU 2301/6-1) to H.B. within the framework of the Priority Programme ‘‘MAdLand – Molecular Adaptation to Land: Plant Evolution to Change’’ (SPP 2237). J.d.V. further thanks the European Research Council for funding under the European Union’s Horizon 2020 research and innovation programme (Grant Agreement No. 852725; ERC-StG ‘‘TerreStriAL’’).

## AUTHOR CONTRIBUTIONS

Conceptualization, J.d.V.; investigation, M.J.B., T.D., S.d.V., H.B., T.P., I.I., J.d.V.; writing – original draft, M.J.B., T.D., S.d.V., T.P., and J.d.V.; writing – review and editing, all authors; visualization, M.J.B., T.D., J.M.R.F.-J., T.P., and J.d.V.; and funding acquisition, J.d.V.

## METHODS

### RESOURCE TABLE

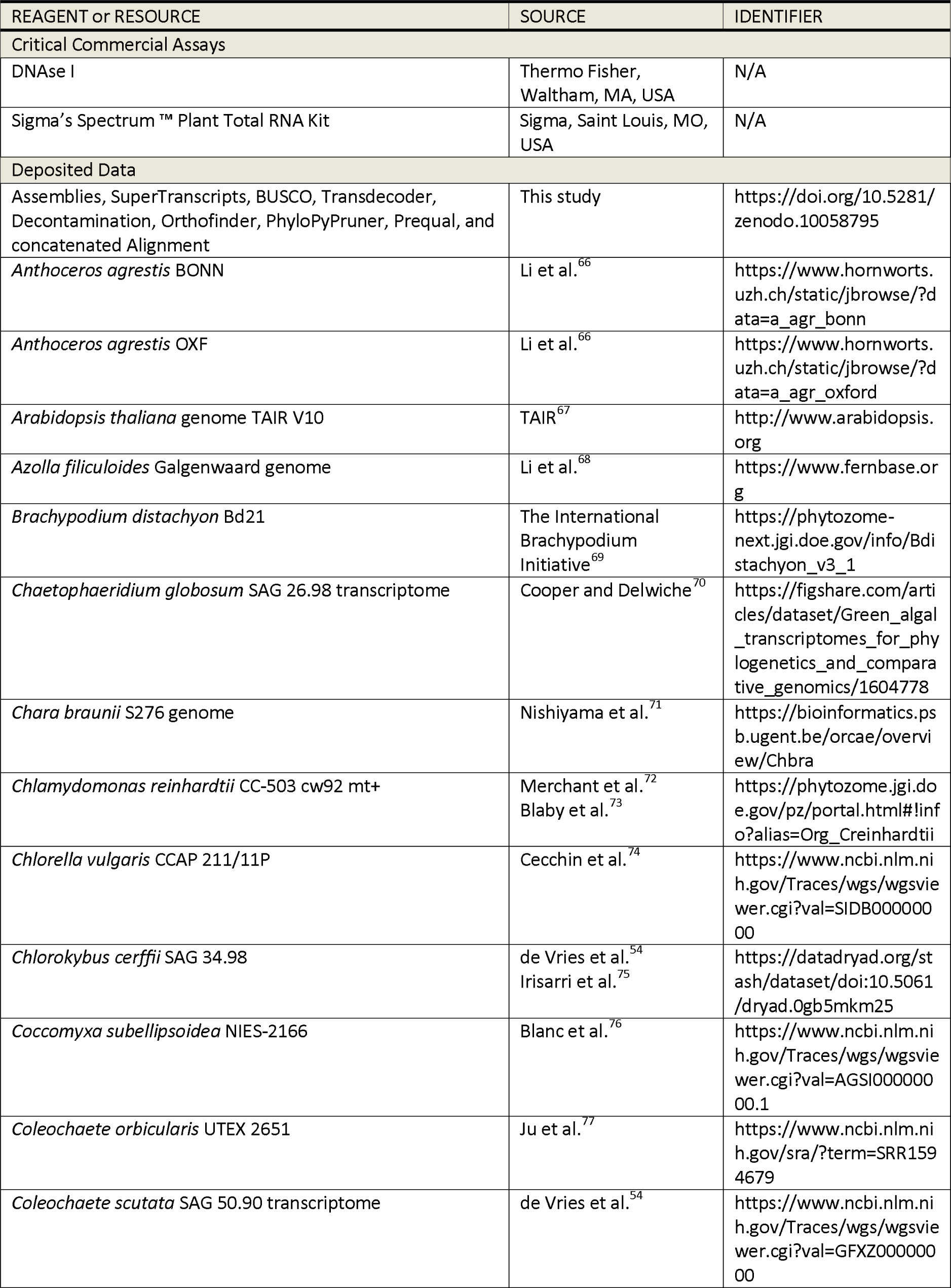
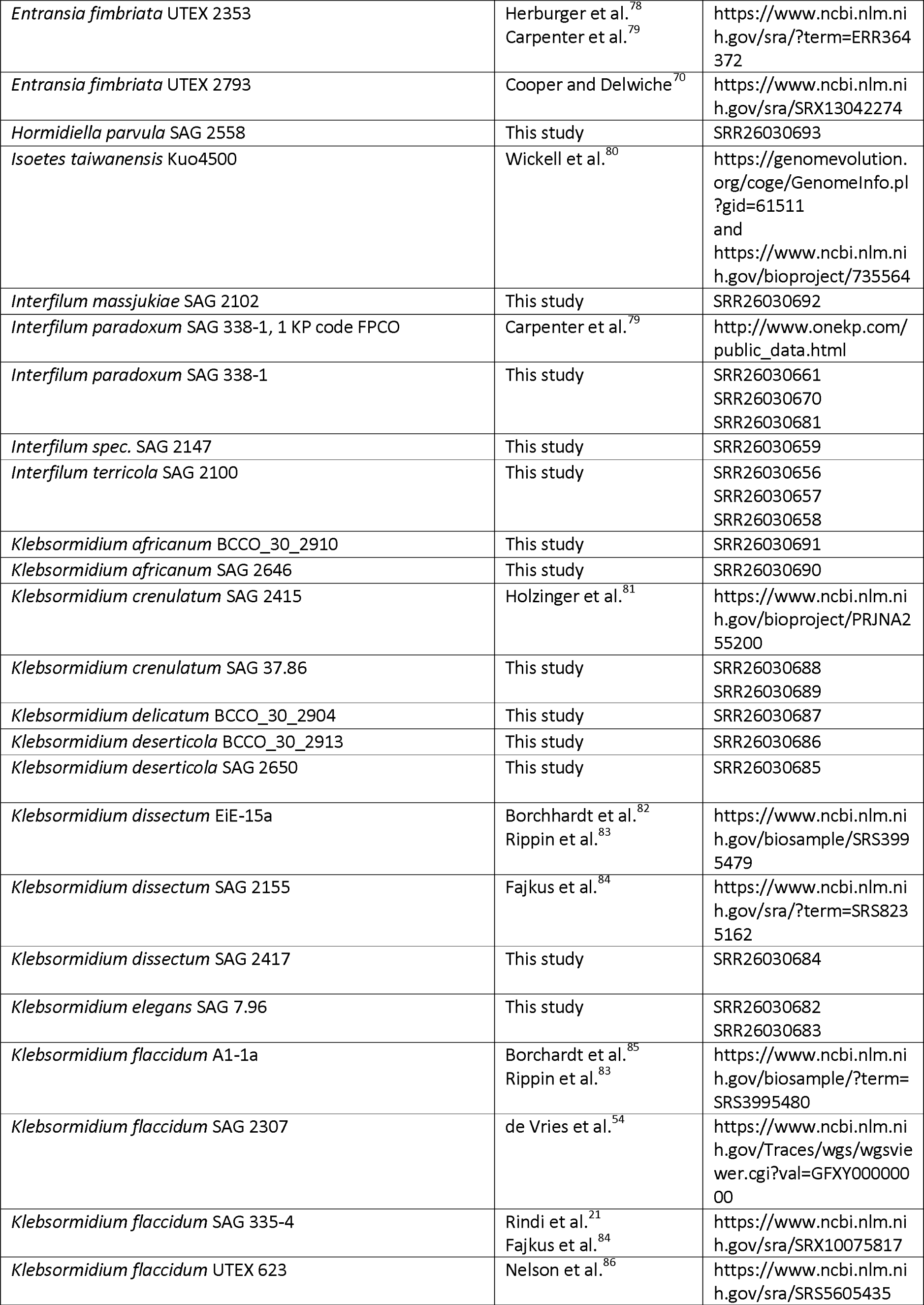
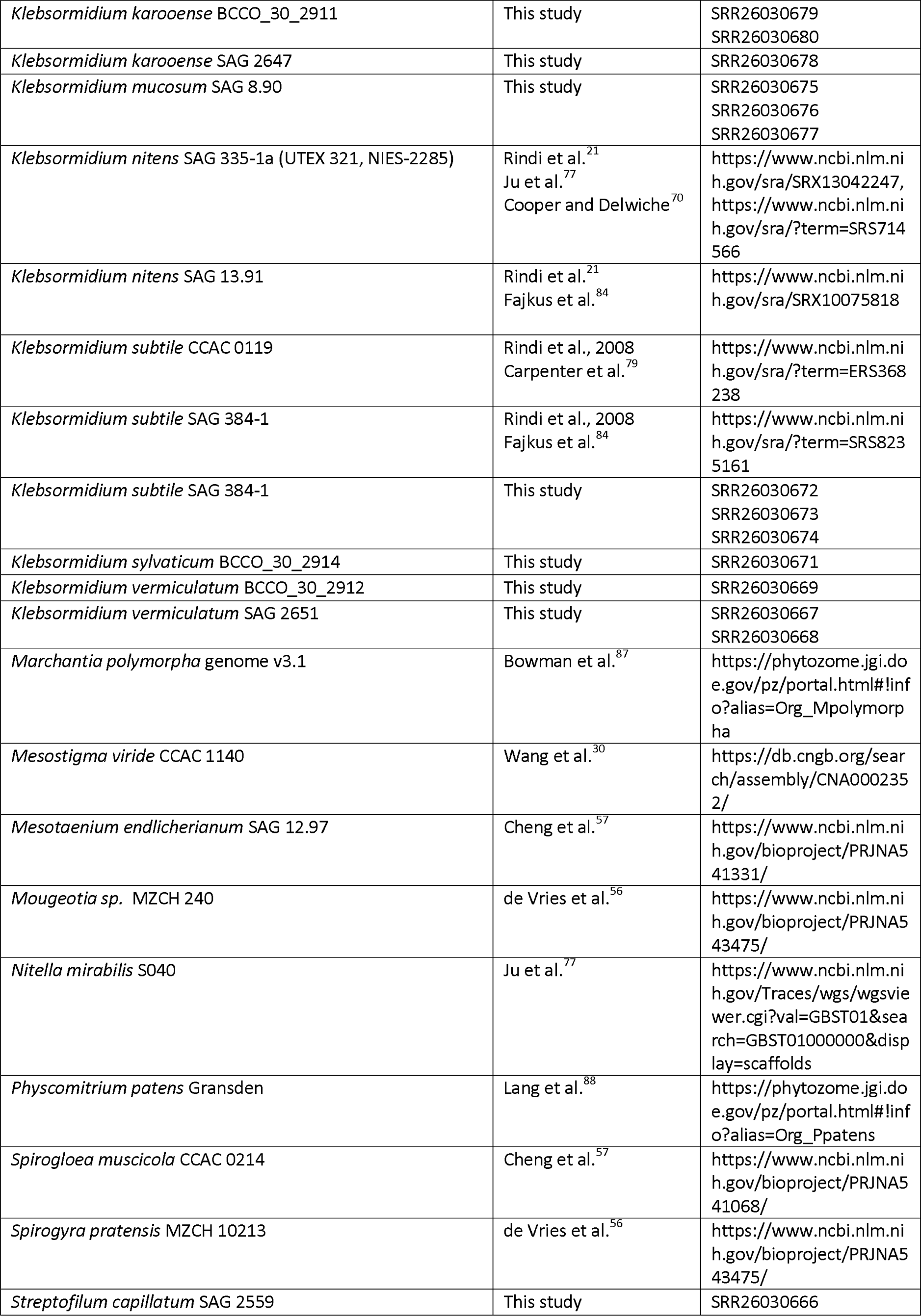
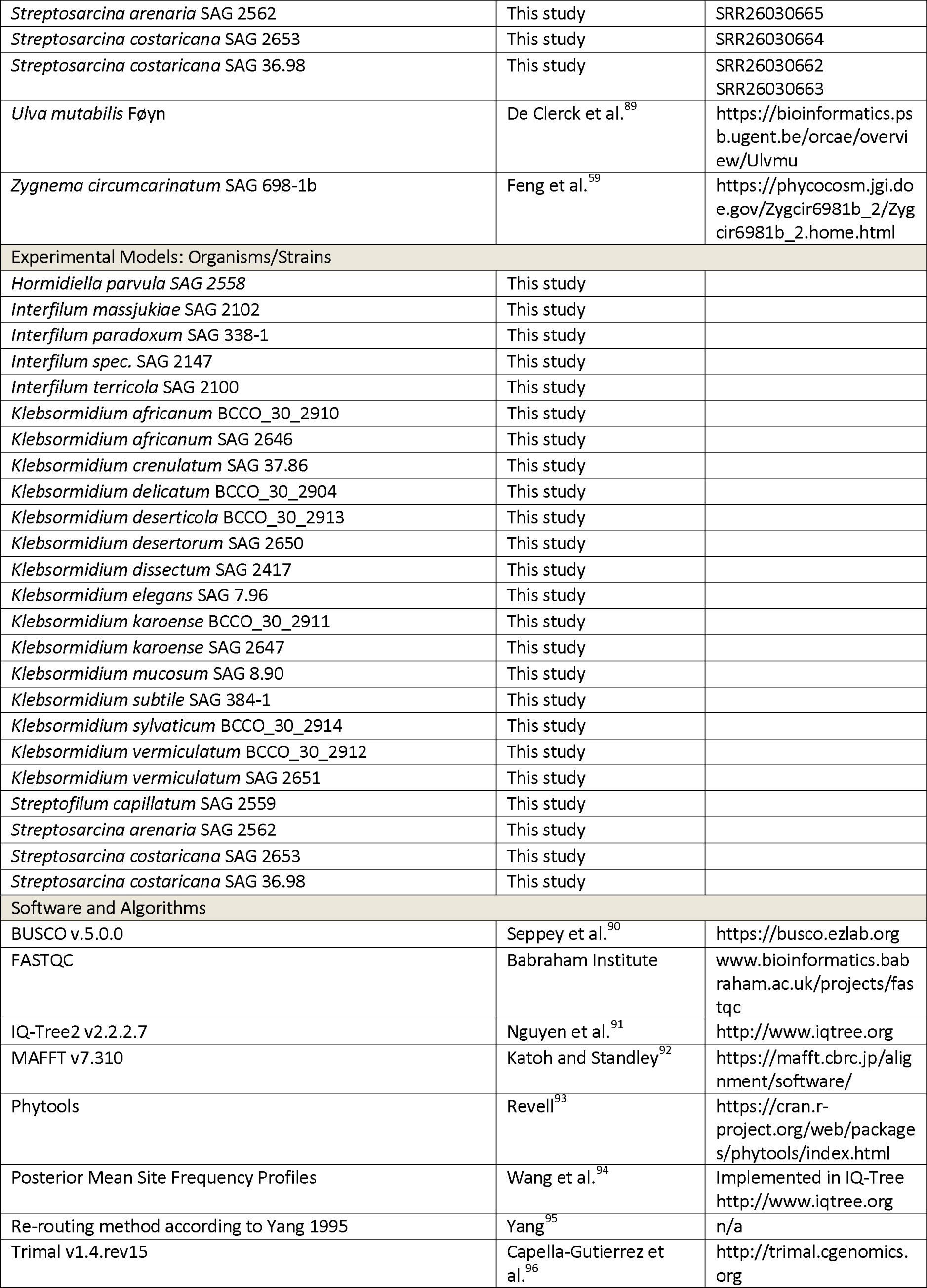
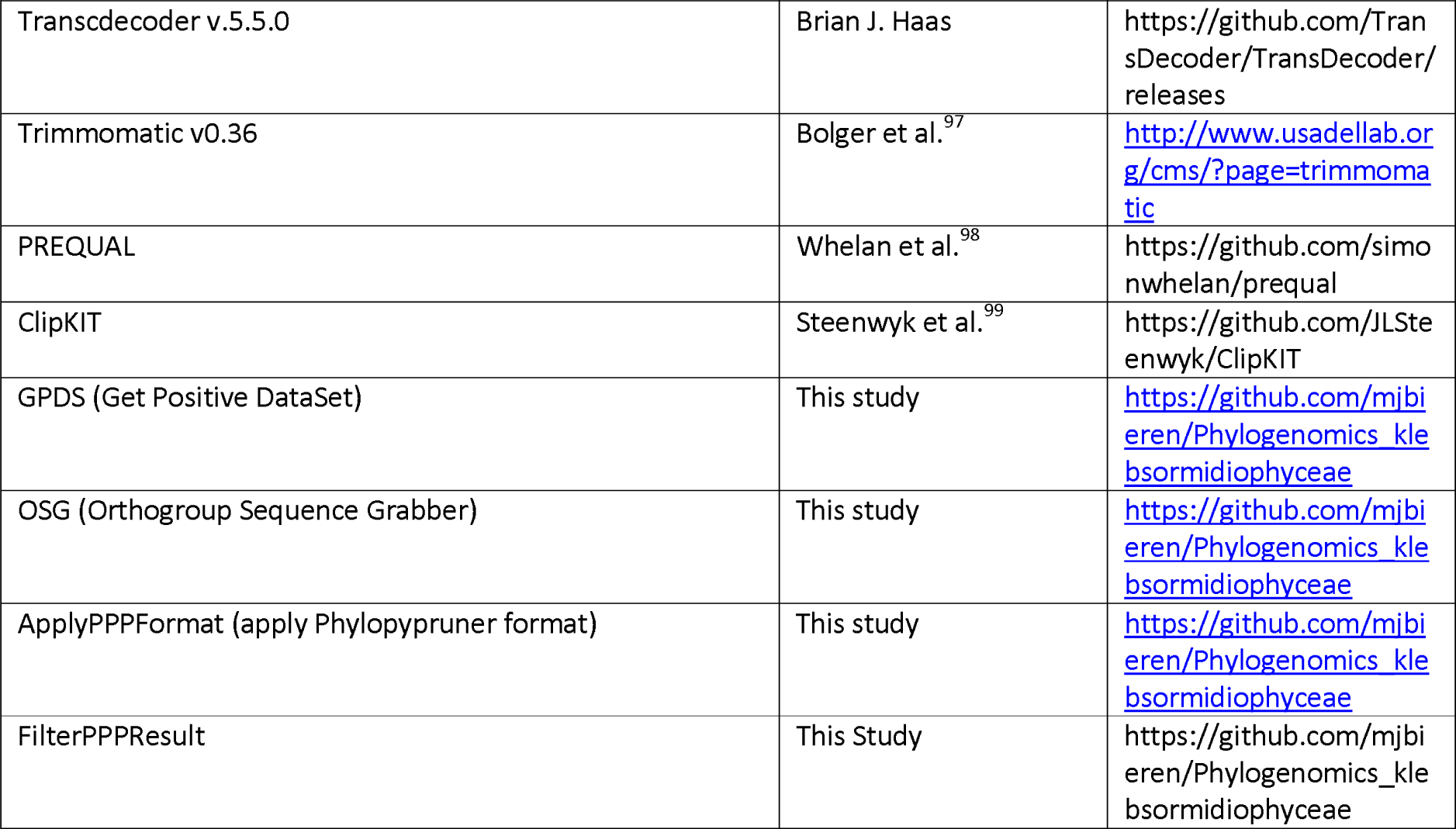

### RESOURCE AVAILABILITY

#### Lead contact

Further information and requests for resources and reagents should be directed to and will be fulfilled by the lead contact, Jan de Vries.

#### Materials availability statement

This study did not generate new unique reagents.

#### Data and code availability statement

- RNA-seq data have been deposited at the NCBI under the BioProject accession PRJNA1013714 and the Sequence Read Archive (SRA) under the accessions SRR26030656-SRR26030693; all data are publicly available as of the date of publication. Accession numbers are additionally listed in the key resources table.
- RNA-seq FastQC reports, BUSCO scorings, transcriptome assemblies, supertranscripts,TransDecoder outputs, decontaminated fasta files, preliminary orthogroups, Phylopypruner result, Prequal; ginsi; clipkit result, the concatenated alignment file and the tree file can be found on Zenodo doi: 10.5281/zenodo.10058795
- The source code for the novel tools discussed in the paper (See RESOURCE TABLE - Software and Algorithms) are available on the GitHub link https://github.com/mjbieren/Phylogenomics_klebsormidiophyceae — any additional computational analyses, not pertaining to these tools, were conducted using established software and are properly referenced in the methods section. Corresponding batch and/or python scripts are also found within the GitHub link

### EXPERIMENTAL MODEL AND SUBJECT DETAILS

#### Algal strains

Strains were obtained from the Culture Collection of Algae at Göttingen University^100^ (SAG). Six authentic strains representing the recently described species of *Klebsormidium*^28^ were received from one of the authors (AL) and officially deposited in at the Culture collection at Institute of Soil Biology (BCCO), Ceske Budejovice, Czech Republic. All strains were cultivated in 3NBBM (medium 26a^101^) at 18°C under full-spectrum fluorescent lamps (25-35 µmol photons m^−2^ s^−1^; 14:10h light-dark cycle).

#### Light microscopy

High-resolution images of the studied strains were done with Olympus BX-60 microscope (Olympus, Japan) with DIC equipped with a ProgRes C14plus camera and the ProgRes® CapturePro Software (version 2.9.01) (JENOPTIK AG, Jena, Germany). All investigated strains examined at the 21st day of cultivation.

#### RNA isolation

For the RNA extraction of 24 different strains, 50 ml of 21-day old liquid culture were centrifuged for 5 min at 20 °C and 11000 rpm and the supernatant were removed. The pellet was transferred into the Tenbroek tissue homogenizer and each sample was manually disrupted during 10 min on ice. Further extraction was done using RNA the Spectrum™ Plant Total RNA Kit (Sigma-Aldrich Chemie GmbH, Germany) according to the manufacturer’s instructions. DNAse I treatment (Thermo Fisher, Waltham, MA, USA) was applied to the RNA samples, and their quality and quantity were assessed using a 1% agarose gel with a SDS stain, and nanodrop (Thermo Fisher), respectively. The RNA samples were shipped on dry ice to Novogene (Cambridge, UK).

#### RNAseq and transcriptome assembly

At Novogene (Cambridge, UK), the samples underwent quality checks using a Bionanalyzer (Agilent Technologies Inc., Santa Clara, CA, USA), and library preparation was performed based on polyA enrichment and using directional mRNA library preparation. The libraries were quality checked and sequenced using the NovaSeq 6000 platform (Illumina) with Novogene dual adapters: 5’- AGATCGGAAGAGCGTCGTGTAGGGAAAGAGTGTAGATCTCGGTGGTCGCCGTATCATT-3’ for read 1 and 5’- GATCGGAAGAGCACACGTCTGAACTCCAGTCACGGATGACTATCTCGTATGCCGTCTTCTGCTTG-3’

Additionally, we downloaded RNAseq data for 14 different Klebsormidiophyceae species, including *Entransia fimbriata* UTEX LB 2353 (ERS368240; Carpenter et al.^79^), *Entransia fimriata* UTEX 2793 (SRR16849194), *Interfilum paradoxum* SAG 338-1 (ERS1830152; Carpenter et al.^79^), *Klebsormidium crenulatum* SAG 2415 (SRS693696, SRS693678, SRS693690; Holzinger et al.^81^), *Klebsormidium dissectum* EiE-15a (SRS3995479; Borchhardt et al.^82^; Rippin et al.^83^), *Klebsormidium flaccidum* A1-1a (SRS3995480; Borchhardt et al.^85^; Rippin et al.^83^), *Klebsormidium flaccidum* SAG 2307 (SRP115828, SRP116582; de Vries et al.^54^), *Klebsormidium flaccidum* SAG 335-4 (SRS8235163; Fajkus et al.^84^), *Klebsormidium flaccidum* UTEX 623 (SRS5605435; Nelson et al.^86^)*, Klebsormidium nitens* SAG 2155 (SRP305831; Fajkus et al.^84^), *Klebsormidium nitens* SAG 13.91 (SRS8235164; Fajkus et al.^84^), *Klebsormidium nitens* SAG 335-1a (SRS10979560, SRS714566; Ju et al.^77^; Cooper and Delwiche^70^), *Klebsormidium subtile* CCAC 0119 (ERS368238; Carpenter et al.^79^), and *Klebsormidium subtile* SAG 384-1 (SRS8235161; Fajkus et al.^84^). All samples’ transcriptomes were assembled *de novo* using Trinity v2.11.0 (Haas et al.^102^) after adapter trimming with Trimmomatic^97^ (--trimmomatic “ILLUMINACLIP:/home/uni08/applbioinfdevries/DATA_ILLUMINA/novogene_adapter_sequences.fa:2:30:10:2:keepBothReads LEADING:3 TRAILING:3 MINLEN:36”). SuperTranscript (Davidson et al.^103^) was inferred by collapsing splicing isoforms using the Trinity implementation. The completeness of the transcriptomes was assessed with BUSCO v5.4.3 (Seppey et al.^90^) using the ‘eukaryota_odb10’ reference set. The combined complete BUSCOs of all newly assembled transcriptomes recovered an average of 90.62% (see data on Zenodo, https://doi.org/10.5281/zenodo.10058795). Protein-coding genes were identified using Transdecoder v5.5.0, with *Klebsormidium nitens*^42^ (NIES-2285) as the reference in BLASTP searches, retaining only the longest open reading frame (ORF) per transcript (--single_best_only).

#### Dataset construction for phylotranscriptomics

To remove potential contaminants, we conducted sequence similarity searches against a comprehensive database that included proteins from various sources. These sources include *Klebsormidium nitens* (NIES-2285)^42^, as well as potential contaminants such as RefSeq^104^ representative bacterial genomes (11,318 genomes), fungi (2,397), all available viruses, archaea (1,833), and plastid genes (78,2087). We employed MMseqs2 (Steinegger and Söding^105^) for the search, using an iterative approach with increasing sensitivities and maintaining a maximum of 10 hits (--start-sens 1 --sens-steps 3 -s 7 --alignment-mode 3 --max-seqs 10). To ensure stringent decontamination, we retained with the help of GPDS (See Resource Table - Software and Algorithms) only sequences that showed the best match to predicted Klebsormidiophyceae nuclear proteins for phylogenetic analysis. For each type of contaminant (bacteria, fungi, viruses, archaea, and plastids), separate files were automatically generated, which can be accessed at Zenodo under https://doi.org/10.5281/zenodo.10058795

#### Phylotranscriptomic analysis

To infer orthogroups, Orthofinder v2.5.4 (Emms and Kelly^106^) was employed using a species tree following the approach of Leebens-Mack et al.^2^.The species tree included representation from chlorophytes, *Chlorokybus cerffii* SAG 34.98, *Mesostigma viride* NIES-296, and various Phragmoplastophyta (see data deposited on Zenodo, https://doi.org/10.5281/zenodo.10058795). This tree also included all the Klebsormidiophyceae with unresolved relationships.

From a total of 1,761,660 orthogroups, 16410 were selected through taxonomic group filtering with the help of OSG (See Resource Table - Software and Algorithms). This selection criterion required the presence of at least one sequence from each of the 10 different taxonomic groups out of the total 14. Additional information regarding the taxonomic group ordering can be found on Zenodo.

Homologous sets were aligned using MAFFT^92^ v7.304 with default settings, and maximum-likelihood inference was performed using IQ-Tree2^91,107^ multicore version 2.2.2.7. The analysis involved fast searches, BIC-selected best-fit nuclear models, and SH-like aLRT branch support (-fast -st AA -m TEST -msub nuclear -alrt 1000). The resulting tree files were transformed into a format compatible with Phylopypruner v1.2.4, as detailed by Thalen et al. (https://pypi.org/project/phylopypruner/), using the assistance of ApplyPPPFilter (see Resource table – Sotware and Algorithms).

Orthologue sets were pruned using Phylopypruner v1.2.4 (Thalen et al., https://pypi.org/project/phylopypruner/) to remove paralogs (--mask pdist --prune MI --min-taxa 10 --trim-lb 5 --min-support 0.75 --min-gene-occupancy 0.1 --min-otu-occupancy 0.1 --threads 80 --trim-freq-paralogs 4 --trim-divergent 1.25 --min-pdist 1e-8 –jackknife), resulting in a set of 5290 orthologues.

After applying the taxonomic filter using FilterPPPResult (-t 3) (see Resource table – Sotware and Algorithms), we identified and selected 2,258 loci. These loci underwent masking with PREQUAL^98^ v1.02. Following this step, we aligned them using MAFFT^92^ ginsi v7.304b with the utilization of a variable scoring matrix (’--allowshift --unalignlevel 0.8’), and any columns containing over 75% gaps were subsequently eliminated using ClipKIT^99^ v2.0.1.

The resultant trimmed alignments were then combined into a matrix consisting of 62 taxa and 420 loci. This matrix comprised a total of 141,000 aligned amino acid positions. To perform maximum-likelihood inference, we employed IQ-Tree2^108^, specifically utilizing the multicore version 2.2.2.7.6. Our analysis consisted of rapid searches, the selection of best-fit nuclear models based on the Bayesian Information Criterion (BIC), and SH-like aLRT branch support (-fast -st AA -m TEST -msub nuclear -alrt 1000). The final tree was then reconstructed using the concatenated file and its partition file using IQ-Tree2 v2.2.2.7 (-m TEST -msub nuclear -s concatenated.fas -p partition.txt -bb 1000 -alrt 1000)

#### Ancestral character state reconstruction

To infer marginal ancestral state estimates for internal nodes in the tree, we conducted ancestral character state reconstruction (ACSR) using Phytools^93^. This software implements Yang’s re-rooting method^95^. We performed two separate ACSR analyses, considering different character coding schemes, to examine the impact on the inferred ancestral character states. The first analysis utilized a 2-character state model, distinguishing between (1) unicellular and (2) multicellular *sensu lato* (including filamentous or multicellular forms). The second analysis employed a 4-character state model, differentiating between (1) unicellular, (2) coccoid, (3) filamentous, and (4) multicellular *sensu stricto*. In all models, we assumed unordered states with equal rates of change.

#### Molecular clock

Bayesian molecular dating was performed with MCMCTree^109^ within the PAML^110^ package v4.9h. We used the phylotranscriptomic tree and eight fossil calibrations with uniform prior distributions (split between Chlorophytes and Streptophytes, Streptophyte crown group and six calibrations within land plants), following parameterizations in Morris et al.^31^ (their Supplementary Table 8). CorrTest^111^ rejected the independent rates model on the maximum likelihood tree (score = 0.89275; p<0.01). Thus, we assumed relaxed autocorrelated lognormal molecular clocks (clock = 2) and birth-death tree priors. Analyses used approximate likelihood calculations^112^ on the phylotranscriptomic dataset (single partition) under the LG+Γ model. A diffuse gamma Dirichlet prior was used for the prior on mean rates as 0.1426 replacements site^−1^ 10^8^ Myr^−1^ (‘rgene_gamma’; α = 2, β = 14.03). The rate drift parameter reflected considerable rate heterogeneity across lineages (‘sigma2_gamma’; α = 2, β = 2). A 100 Ma time unit was assumed. Two independent MCMC chains were run for each analysis, consisting of 22 million generations and the first 2,000,000 were excluded as burnin. Convergence was checked using Tracer^113^ v1.7.1; all parameters reached effective sample size (ESS) > 200.

