## Supplemental Figures for "Phylogenomic insights into the first multicellular streptophyte"

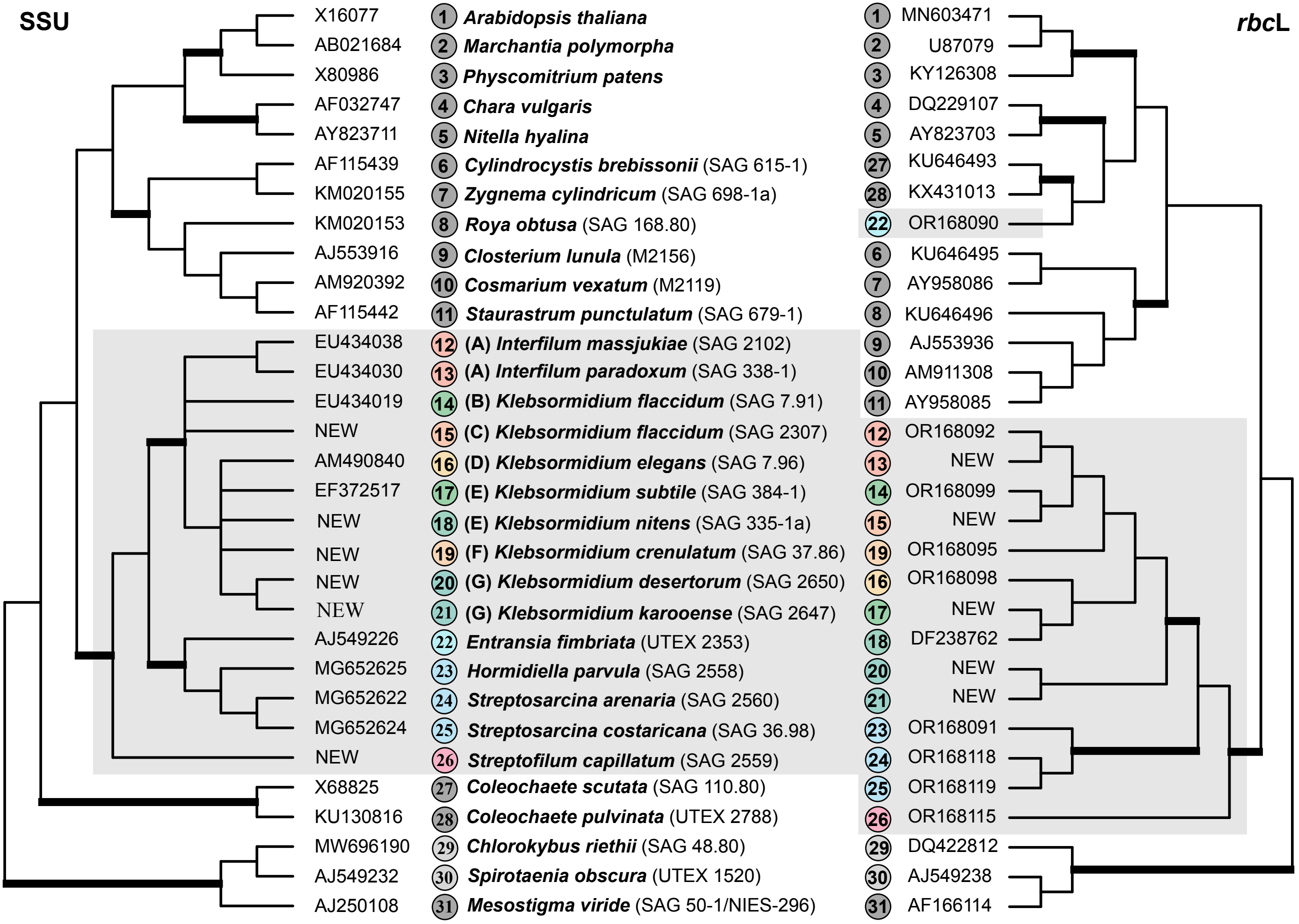


**Figure S1: Molecular phylogeny of the Streptophyta based on SSU rDNA and *rbc*L sequence comparisons, related to Figure 2.** The phylogenetic trees shown were inferred using the maximum likelihood method based on the data sets (31 taxa: 1794 aligned positions for SSU, 1428 for *rbc*L) using PAUP 4.0a169 (Swofford 2002). For the analyses the best model was calculated by the automated model selection tool implemented in PAUP. The setting of the best model was given as follows: (SSU) TrN+I+G (base frequencies: A 0.2453, C 0.2156, G 0.2823, T 0.2569; rate matrix A-C 1, A-G 2.0336, A-U 1, C-G 1, C-U 6.2892, G-U 1) with the proportion of invariable sites (I = 0.4778) and gamma shape parameter (G = 0.5542); (*rbc*L) GTR+I+G (base frequencies: A 0.2913, C 0.1835, G 0.1859, T 0.3392; rate matrix A-C 1.1839, A-G 6.1340, A-U 3.4723, C-G 1.2456, C-U 9.5079, G-U 1.0000) with the proportion of invariable sites (I = 0.5316) and gamma shape parameter (G = 2.222). The branches in bold are highly supported in all analyses (bootstrap values > 70% calculated with PAUP using maximum likelihood, neighbor-joining using maximum likelihoodsettings, and maximum parsimony). The classes are marked in the same colors as in the main Figure 2.

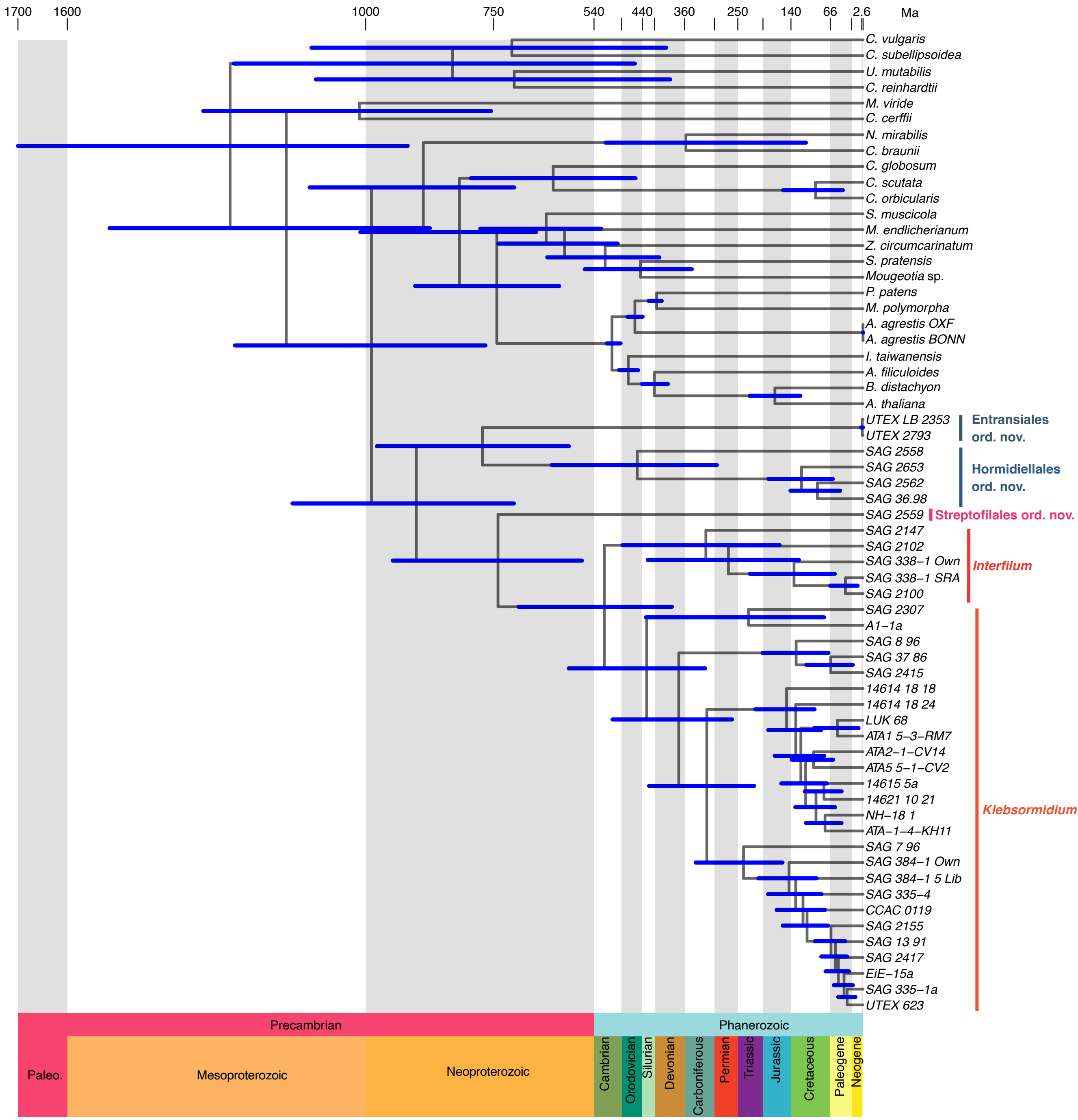


**Figure S2: Molecular clock analysis of the Klebsormidiophyceae, related to Figures 2 and 3.** Estimates of divergence times in million years were calculated using Time Tree. A uniform distribution was applied. Chlorophytes served as outgroup.

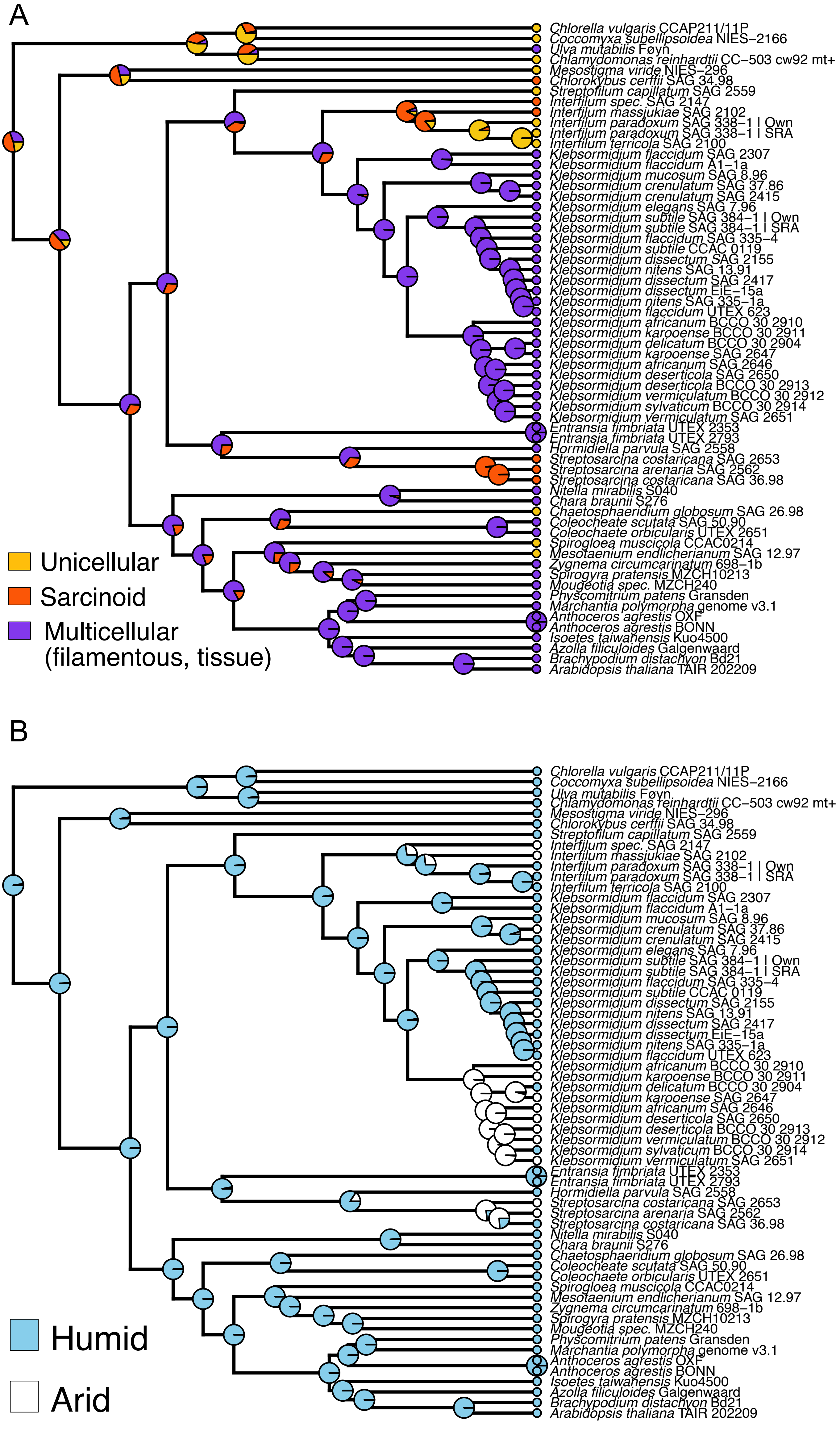


**Figure S3: Ancestral character state reconstruction of body plan across 800 million years of klebsormidiophyceaen evolution, related to Figure 3. A**: To examine the ancestral character states of growth types in unicellular or multicellular organisms, coding schemes represented varying levels of complexity and hypotheses regarding the homology of growth types. The shown color-coded character state distributions represents yellow for unicellular growth, orange for sarcinoid growth, purple for filamentous growth, and purple for multicellular growth sensu stricto. **B**: To examine the ancestral habitats of the Klebsormiodphyceae, we coded the habitat occurrence of the species as light blue for humid and white for arid. Note the arid-dwelling ancestor of the G-clade of *Klebsormidium* spp.

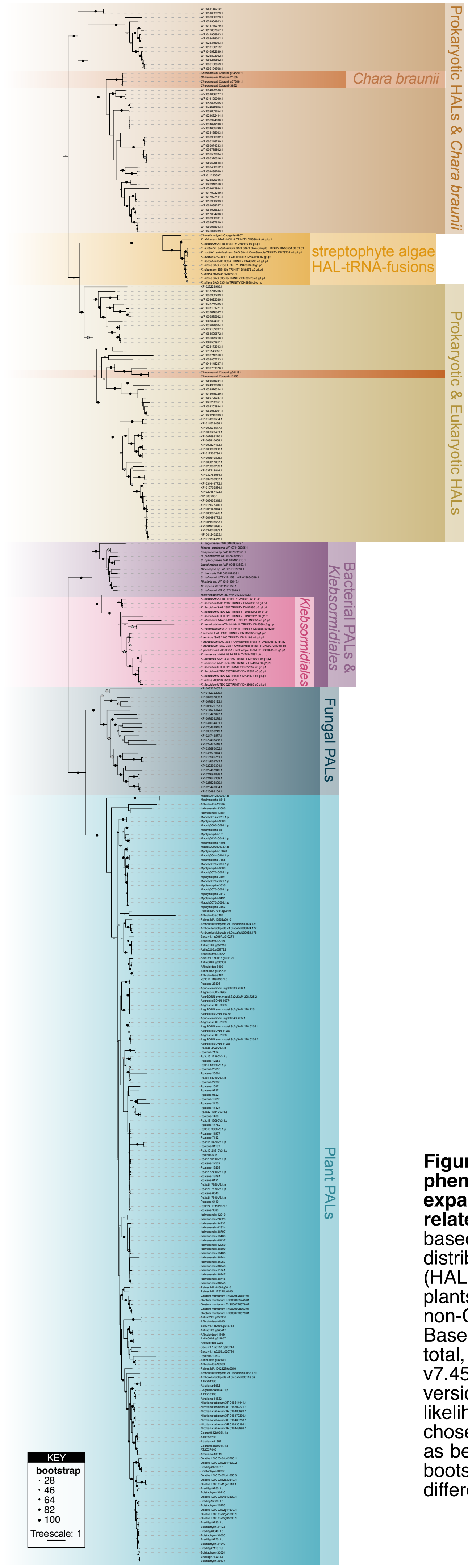


**Figure S4: Phylogenetic analysis of phenylalanine ammonia-lyase (PAL) with an expanded sampling in Klebsormidiophyceae, related to Figure 3.** Sequences were sampled based on de Vries et al. (2021), with a phylodiverse distribution of PAL and histidine ammonia lyase (HAL) sequences from (i) the green lineage (land plants, streptophyte algae and chlorophytes), (ii) non-Chloroplastidial eukaryotes, and (iii) bacteria. Based on these 364 PAL and HAL sequences in total, an alignment was computed using MAFFT v7.453 with a L-INS-I approach. IQ-TREE multicore version 1.5.5 was used to compute a maximum likelihood phylogeny with LG+F+G4, which was chosen according to Bayesian Information Criterion as best-fit-model using ModelFinder. 5000 ultrafast bootstrap replicates were computed (shown in differently sized dots, see KEY).

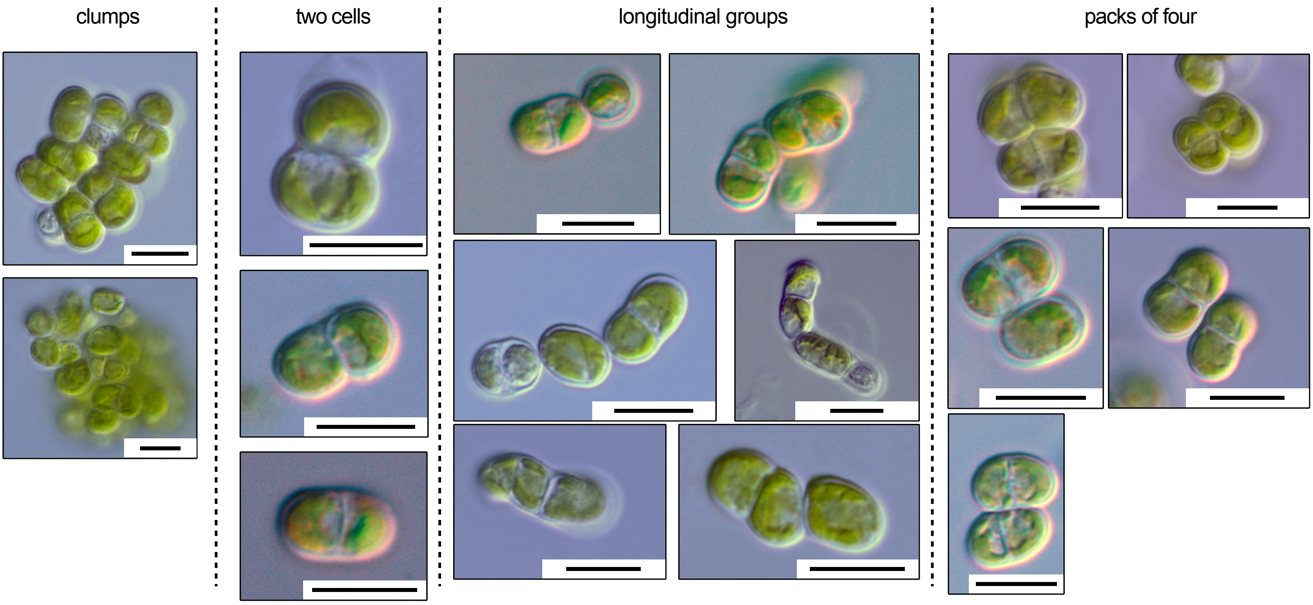


**Figure S5: Different morphologies of *interfilum*, related to Figure 3.** *Interfilum* SAG2147 was grown on agar and diverse morphologies were observed using light microscopy. Scale bar = 10 $\mu$m in all pictures.
